# CD96-mediated internalisation of ligand CD155 as a novel mechanism for immune regulation

**DOI:** 10.64898/2026.05.29.727853

**Authors:** Diana Shinko, Rosemarie A Ford, Maja Kos, Meryl H Attrill, Rafter Wu, Reemon Spector, Anne M Pesenacker

## Abstract

CD96 is a member of the immunoglobulin superfamily including TIGIT and CD226 that bind to a shared ligand, CD155. This family of co-receptors and its ligands are dysregulated in various autoimmune conditions and cancers, highlighting therapeutic potential. While TIGIT and CD226 are recognised co-inhibitory and co-stimulatory receptors respectively, the function of CD96 remains incompletely defined. Here, we assessed CD96-CD155 interaction and downstream trafficking to define a novel CD96 mechanism of action. Using primary human T cells and engineered cell models, we demonstrate that CD96 mediates uptake and internalisation of both soluble and cell-associated CD155. CRISPR-Cas9-mediated receptor knockout in primary human T cells revealed that CD155 uptake was uniquely dependent on CD96, but not TIGIT or CD226, identifying CD96 as the dominant mediator of CD155 internalisation in human T cells. This uptake process was dependent on active receptor cycling, which was partially facilitated by the CD96 cytoplasmic domain. Furthermore, CD96 variant 2 exhibited enhanced ligand binding and uptake efficiency compared with variant 1. Mechanistically, internalised CD155 trafficked to lysosomal compartments and associated with autophagy-related proteins, consistent with degradative processing. These findings reveal a previously unrecognised mechanism by which CD96 may regulate ligand availability through ligand internalisation and trafficking, with potential implications for T cell function and immune regulation. This process may promote changes in the immunoregulatory balance and could inform new targeted therapeutic development.

## Introduction

CD96, also known as TACTILE, is a type I transmembrane glycoprotein and a member of the immunoglobulin (Ig) superfamily, which includes TIGIT and CD226, binding nectin and nectin-like proteins^1^. The common ligand, CD155, also known as poliovirus receptor (PVR) or Necl5, is shared between all three co-receptors^2^. This relatively novel co-receptor family has attracted interest for potential therapeutic applications such as in cancer immunotherapy^3, 4^.

CD96 has been reportedly elevated in immune-mediated conditions such as autoimmune vasculitis, poor-prognosis cancers, and in the inflamed joint of individuals with juvenile idiopathic arthritis^5–7^. CD155 upregulation has also been reported in multiple human malignancies, such as colorectal cancer and lung adenocarcinomas, where it is often associated with poor prognosis^2, 8, 9^.

TIGIT is widely considered a co-inhibitory receptor, whereas CD226 functions as a co-stimulatory receptor^3^. TIGIT has been shown to suppress anti-tumour immunity; for instance, studies using pre-clinical mouse tumour models demonstrated that TIGIT-deficient mice exhibited improved overall survival as well as reduced lung metastasis of B16 cells^10, 11^. In contrast, the CD226-CD155 axis has been shown to promote anti-tumour immunity, with CD226 deficiency impairing the cytotoxic and proliferative capacity of CD8+ T cells and NK cells against CD155-expressing tumour cells, resulting in increased tumour burden in preclinical mouse models^11–15^.

However, the role of CD96 in immune response is less clearly defined. Some studies suggest that CD96 blockade promotes NK and CD8+ T cell anti-tumour activity in mice suggesting co-inhibitory action^16, 17^. Similarly, a role for CD96 has been reported in controlling inflammation whereby CD96KO mice exhibit elevated NK cell IFN-y production and hyperinflammatory responses upon LPS challenge^15^. In contrast, other literature suggests that CD96 cross-linking could enhance human NK and CD8+ T cell activation^7,8^, and that T-cell specific CD96-deficiency may produce milder symptoms in murine psoriasis models^18^. Similarly, others demonstrate CD96 engagement potentially driving IFN-y production in addition to pro-inflammatory cytokines^19^. Therefore, CD96 function is still poorly understood and needs further investigation to explain these conflicting findings.

In addition to direct signalling^20^ and competitive binding for ligand^21, 22^, co-receptors may also regulate immune responses by modulating ligand availability^23^. This process has been well described in the CTLA4/CD28 checkpoint axis, where CTLA-4 mediates transendocytosis of CD80/CD86 removing the ligands from the environment, reducing their availability to CD28 and thus decreasing immune activation^23, 24^.

In this study, we assessed CD96-CD155 interaction and downstream trafficking to define a novel CD96 mechanism of action. We show that CD96-CD155 ligation promotes internalisation and degradation, supporting a model in which CD96 modulates ligand availability. This process may regulate CD155 availability to TIGIT and CD226, thereby influencing their opposing effects on T and NK cell activation. Therefore, our data provides a potential explanation for previously conflicting reports on CD96 function. These findings will be critical in understanding this novel co-receptor family to inform new treatment development targeting these co-receptors across immune-mediated conditions such as cancer and inflammation.

## Results

### CD96 is highly expressed on T cells

To assess the expression of CD96 on human immune cell types, we utilised our previously published spectral flow cytometry data from the peripheral blood of healthy donors (n=18, FLOWRepository ID FR-FCM-Z6VC)^6^. Our results showed that B cells had the lowest expression of CD96 (8.01-2214 min-max MFI), myeloid cells such as classical dendritic cells had a high but variable expression (786.0-8056.0 MFI), and T cells had a consistently high expression of CD96 (1908.0-4934.0 MFI, **Figure 1A, B**).

**Figure 1.**
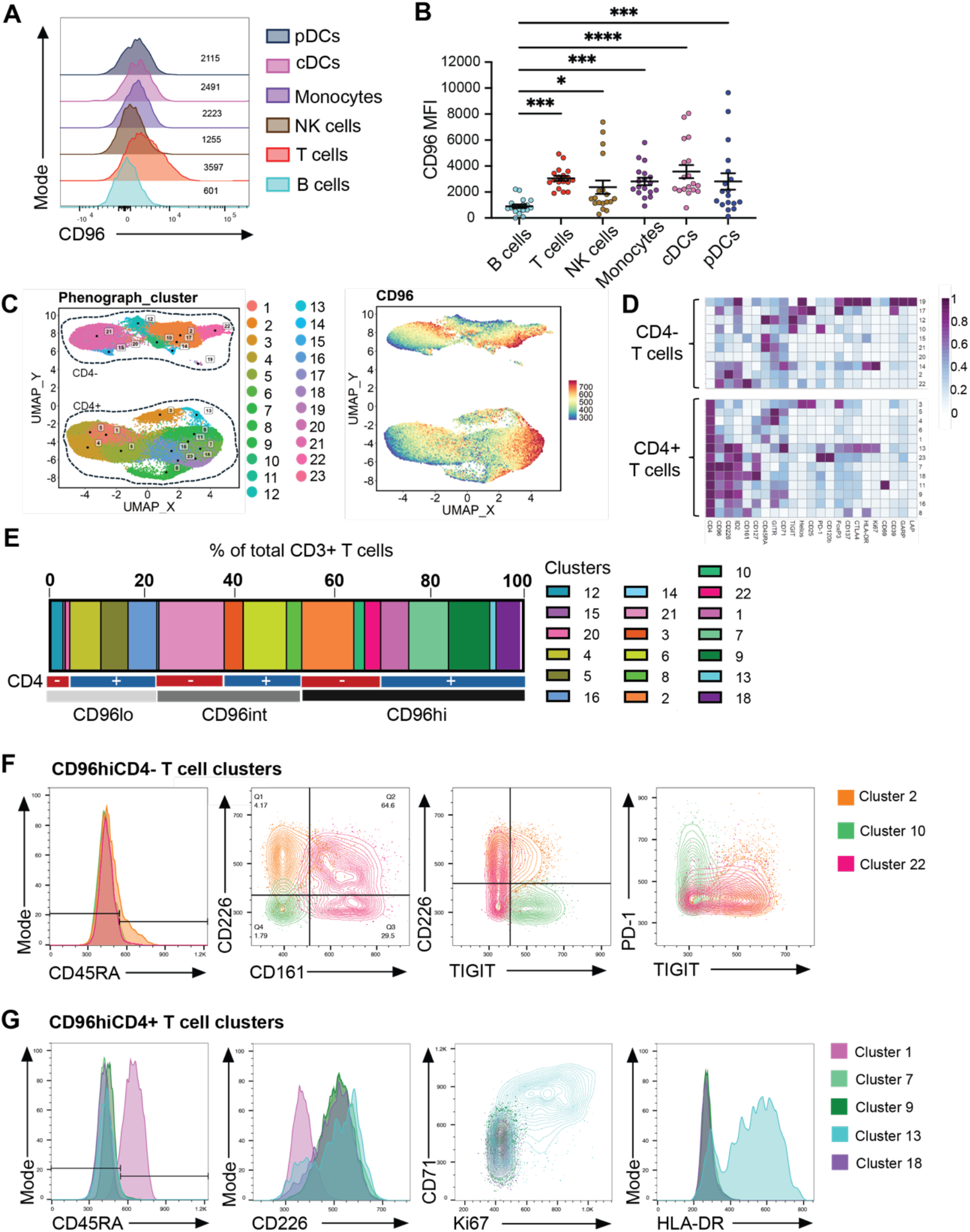
CD96 is highly expressed on T cells. Spectral flow cytometry data of healthy control PBMC (n=18) from Attrill et al, 2024 was analysed. **(A)** Representative histogram and **(B)** median fluorescence intensity (MFI) of CD96 in various immune cell subsets using manual gating. Data as mean±SEM and two-way ANOVA with Tukey’s multiple comparisons test. **(C)** Unbiased clustering algorithm PhenoGraph, after gating on all single, live (FVD−), CD3+CD19-CD56-cells, identified 23 clusters of T cells with UMAP of all samples, and with CD96 MFI overlayed (right). **(D)** T cell markers heatmap of CD4- and CD4+ T cells clusters, Z-score across columns. **(E)** Frequency of T cell clusters (as a proportion of total CD3+ cells), stratified by CD96 MFI expression level (low (lo), intermediate (int), and high (hi)) and CD4 expression. Overlay flow plots differentiating **(F)** CD96hiCD4-T cell clusters (2, 10, 22) and **(G)** CD96hiCD4+ T cell clusters (1, 7, 9, 13, 18). *p<0.05, ***p<0.001, ****p<0.0001.

To further characterise CD96 expression across *ex vivo* T cell subsets, CD3+CD19-CD56-cells were clustered unbiasedly by PhenoGraph^25, 26^ (**Figure 1C-D**). The clustering algorithm produced 23 clusters, with clusters with a frequency <0.25% excluded from further analysis resulting in 8 CD4- and 11 CD4+ clusters assessed for CD96 expression and phenotype. In total, CD96 intermediate or high clusters made up 76.47% of total CD3+ T cells (**Figure 1E**), demonstrating the high prevalence of CD96 across T cell subsets. CD4-CD96hi clusters were predominantly memory (CD45RA-) and co-expressed CD161, CD226, TIGIT, and/or PD-1 (**Figure 1D, F**). CD4+ CD96hi clusters comprised both naïve and memory populations and including a highly activated and proliferative subset (**Figure 1D, G**). Given the diversity of CD96 expression, the expanding appreciation of co-receptor biology, and the focus of prior literature on the TIGIT/CD226/CD96 axis on CD8+ T cells and NK cells, this work instead focuses on primary CD4+ T cells, where these pathways remain less well characterised, and utilises cell line models to understand CD96 mechanisms.

### CD155 uptake and internalisation by T cells is dependent on CD96

The role of CD96 remains poorly defined. Co-receptors can mediate their effect by direct signalling^20^, competition^21, 22^, and/or modulation of shared ligand availability^23^. First, we investigated CD96 interaction with soluble, recombinant ligand-Fc fusion protein (CD155Fc). Human primary CD4+ T cells were isolated and expanded to maximise CD96 expression (**supplemental Figure S1A**), incubated with CD155Fc, and stained with anti-human Fc antibody either on the surface or post-fixation and permeabilisation showing increased total levels of CD155Fc (surface: 24.16±2.52% vs total: 43.70±4.22%, p=0.0039; MFI surface: 1494±546.1 vs total:4050±1579, p=0.0039, **Figure 2A**). Thus, while some CD155Fc was bound to the surface, a larger proportion of CD155Fc was internalised by CD4+ T cells.

**Figure 2.**
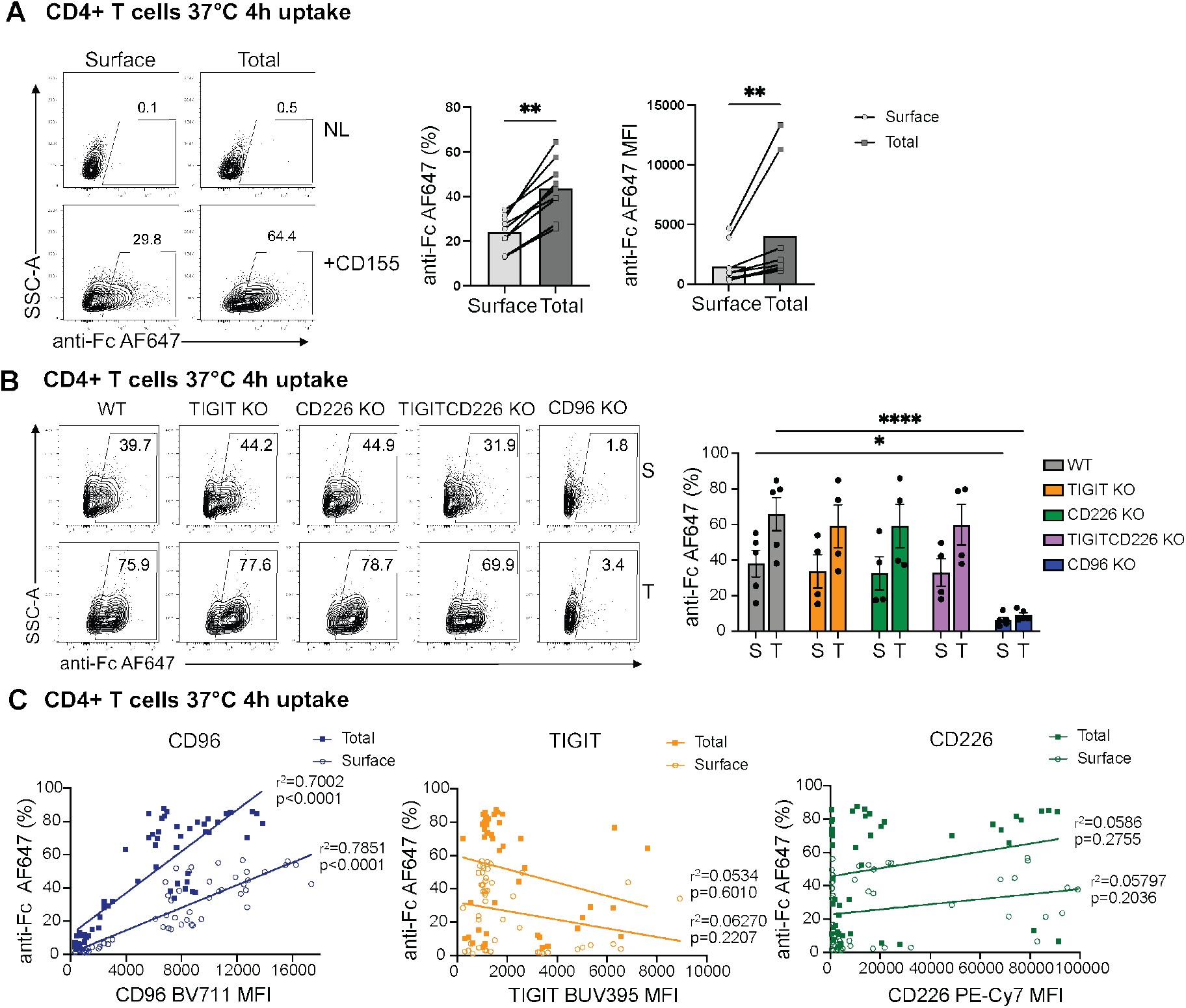
CD155 uptake and internalisation in primary CD4+ T cells is dependent on CD96. Wildtype (WT) or CRISPR-Cas9-mediated knockout (KO) of TIGIT, CD226, TIGIT and CD226, or CD96, expanded, primary human CD4+ T cells were incubated with soluble CD155Fc (5μg/mL) or no ligand (NL) for 4h at 37°C and 5% CO_2_, and stained with anti-human Fc AF647 antibody on the surface or post-fixation/permeabilisation for the total stain. **(A)** Representative plots of WT CD4+ T cells ± ligand (CD155Fc) with the frequency and MFI of anti-human Fc AF647 (CD155) on the surface vs total (n=9). Data analysed using Wilcoxon paired test. **(B)** Representative plots of WT or CRISPRed CD4+ T cells incubated with CD155Fc and frequency of anti-human Fc AF647 (CD155) on the surface vs total (n=5). Data as mean±SEM and two-way ANOVA with Šídák’s multiple comparisons test. **(C)** Correlation between frequency of surface or total anti-human Fc AF647 (CD155) and CD96, TIGIT, or CD226 expression (MFI) on expanded WT or CRISPRed CD4+ T cells, regulatory T cells (Tregs), and conventional T cells (Tconvs). *p<0.05, **p<0.01, ****p<0.0001.

To define whether this ligand uptake was CD96 dependent, we utilised CRISPR-Cas9 to knock out TIGIT, CD226 and/or CD96 (which all share CD155 as ligand and can be co-expressed) in CD4+ T cells/Tregs (**supplemental Figure S1B**). Ligand uptake of CRISPRed cells showed that TIGIT, CD226, or TIGIT CD226 double KO had no effect on CD155Fc staining compared to WT cells (**Figure 2B** representing CD4+ T cells). However, CD96 KO significantly reduced, almost abolished, CD155Fc staining compared to the WT cells (surface: 6.37±1.53% vs 38.10±7.46%, p=0.0316; total: 9.18±1.27% vs 65.84±9.38%, p<0.0001) (**Figure 2B**), indicating that ligand uptake and internalisation was highly dependent on CD96. Furthermore, the expression of CD96 significantly correlated with both surface (r^2^=0.7851, p<0.0001) and total (r^2^=0.7002, p<0.001) CD155Fc staining, whereas TIGIT and CD226 showed no significant correlation (**Figure 2C**). These findings reveal a distinct ability of CD96 to internalise shared ligand CD155.

### CD96 variant 2 is more efficient at binding and uptake of CD155

Human CD96 naturally occurs in 2 variants due to alternate splicing of the pre-mRNA^27^. Evaluating the mRNA gene expression consistently showed a higher expression of CD96 variant 2 (CD96v2) compared to CD96v1 (PBMCs: 4.23±0.10 vs 3.18±0.37; p=0.056, CD4+ T cells 4.69±0.09 vs 3.83±0.18, p=0.0286, Tregs 3.95±0.20 vs 2.70±0.41, p=0.0159, Tconvs 4.60±0.16 vs 3.81±0.26, p=0.0556, **Figure 3A**).

**Figure 3.**
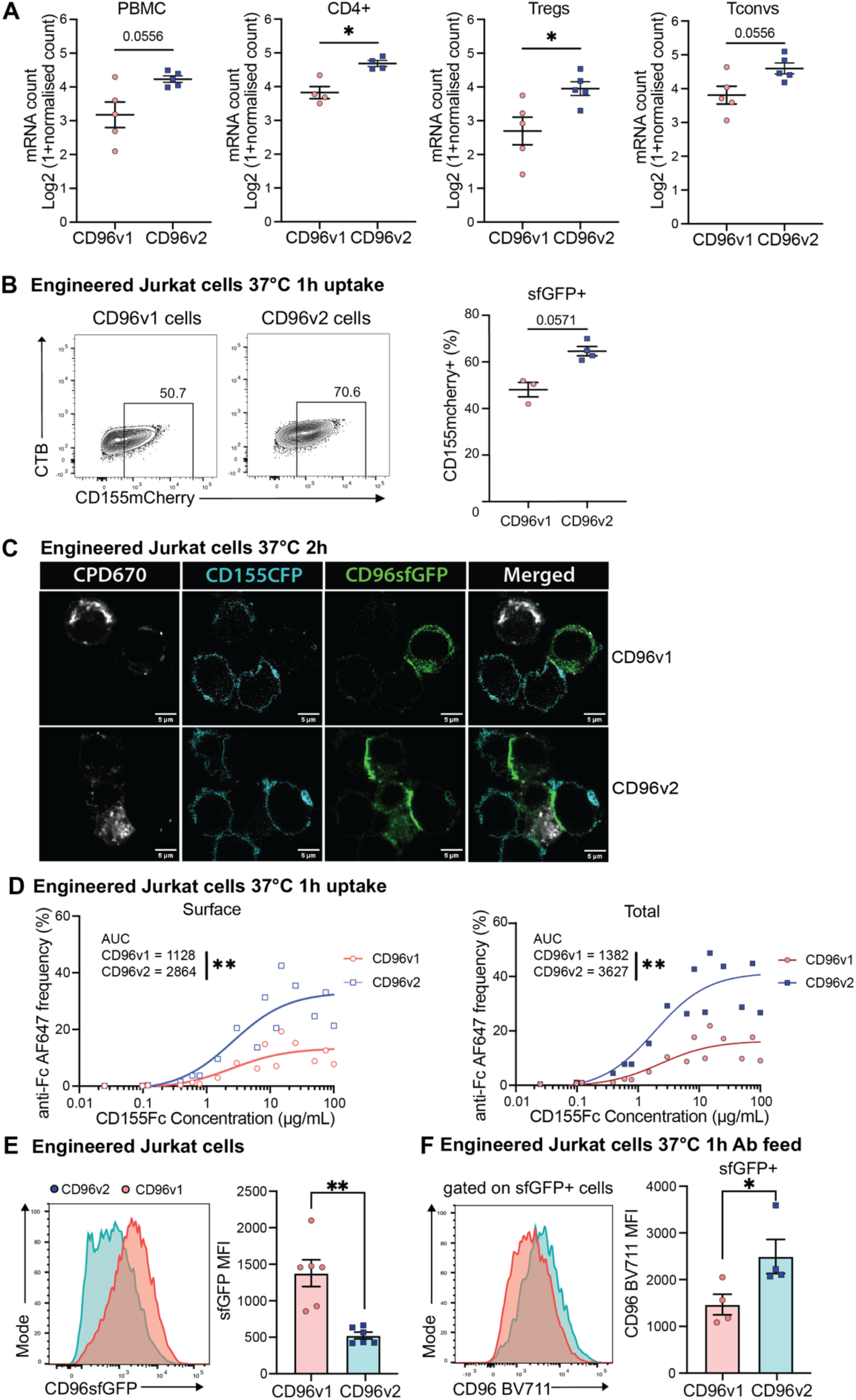
CD96 variant 2 is more efficient at binding and uptake of CD155. **(A)** mRNA count (nanoString) of CD96v1 and v2 gene expression in primary human PBMCs, CD4+ T cells, Tregs, and Tconvs (n=5). Data as mean±SEM and Mann-Whitney test. **(B)** Engineered Jurkat cells expressing CD96v1 or v2sfGFP were incubated 1:1 with DG75.CD155mCherry cells for 1h at 37°C and 5% CO2. Frequency of CD155mCherry in sfGFP+ gated Jurkat cells (n=4). Data as mean±SEM and Mann-Whitney test. **(C)** CPD670-labelled (Cell Proliferation Dye 670, grey) engineered Jurkat cells expressing CD96v1 or v2sfGFP (green) were incubated 1:1 with DG75.CD155CFP (cyan) cells for 2h at 37°C and 5% CO2. Representative confocal microscopy images of the co-cultured cells. Scale bar=5μm. **(D)** Engineered Jurkat cells expressing CD96v1 or v2sfGFP were incubated with soluble CD155Fc at a range of concentrations (0.025-100.0μg/mL) for 1h at 37°C and 5% CO2, stained with anti-human Fc AF647 antibody on the surface or post fixation/permeabilization for the total stain. Frequency of surface and total anti-human Fc AF647 (CD155) (n=2), line represents mean, Welch’s t test comparing AUCs. **(E)** sfGFP levels for engineered Jurkat cells expressing CD96v1/CD96v2-sfGFP fusion protein. Representative histograms (left) and median fluorescence intensity (MFI, right), data as mean±SEM (n=5) and Mann-Whitney test **(F)** Engineered Jurkat cells expressing CD96v1 or v2sfGFP were incubated with anti-CD96 BV711 antibody for antibody feed for 1h at 37°C and 5% CO2. Representative histograms (left) and median fluorescence intensity (MFI, right) of anti-CD96 BV711 staining in sfGFP+ cells (gated based on Ø cells; n=4), Data as mean±SEM and Mann-Whitney test. *p<0.05, **p<0.01.

Work by Meyer et al^27^ suggested that CD96v2 may have higher binding to CD155 compared to variant 1 by (cold) staining with recombinant ligand, but the functional differences between the two variants remain poorly understood. We engineered Jurkat (Jurkat.CD96v2sfGFP, Jurkat.CD96v1sfGFP) and DG75 cells (DG75.CD155mCherry) and showed that in co-culture a higher proportion of CD96v2sfGFP+ cells were CD155mCherry positive compared to CD96v1+ cells (p=0.0571, **Figure 3B**), indicating a greater ability to take up ligand by CD96v2. This may be due to a stronger binding of CD96v2 to CD155 as confocal microscopy imaging showed stronger synapses between CD96v2 and CD155 expressing cells (CFP-tagged CD155 expressing DG75 cells) compared to CD96v1 (**Figure 3C**). Consistent with this, soluble ligand was bound and internalised by more CD96v2 expressing cells than CD96v1 expressing cells in a concentration dependent manner (surface: 2864±337.4 vs 1128±172.8 AUC, p=0.0038; total: 3627±491.6 vs 1382±259.6 AUC p=0.0066, **Figure 3D**). Interestingly, these experiments showed that even though CD96v1sfGFP cells had higher CD96 expression by sfGFP MFI (**Figure 3E, supplemental Figure S2**), CD96v2 cells took up more soluble ligand as well as anti-CD96 BV711 antibody (ligand (**Figure 3D**)/antibody feed at 37°C, gated on sfGFP+ cells: anti-CD96 BV711 MFI 1469.3±220.0 for CD96v1 vs 2497.0±365.8 for CD96v2, p= 0.0286, **Figure 3F**), suggesting both ligand and commercially available antibodies show a preference for CD96v2. Taken together, these results demonstrate that CD96v2 is more prevalent and efficient than CD96v1 in mediating CD155 binding and uptake.

### CD96 cycles and internalises CD155 in an active process

Similar to our engineered Jurkat cells (see **Figure 3**), expanded primary CD4+ T cells showed active cycling in an anti-CD96 antibody feed at 37°C with higher anti-CD96 BV711 MFI than 4°C surface staining (MFI: 5501.0±144.0 vs 2425.0±488.0, **Figure 4A**), with both antibody uptake and cold staining abolished in CD96 KO cells (crCD96 KO gated on CD96-, MFI: 146.0±6.0 vs 206.5±2.5). The antibody feed also indicated continuous active cycling in the cell, as anti-CD96 BV711 MFI increased over time with a peak at 4h. Whether CD96 recycles back to the surface upon internalising or new CD96 is constantly replenished to the cell surface remains to be seen.

**Figure 4.**
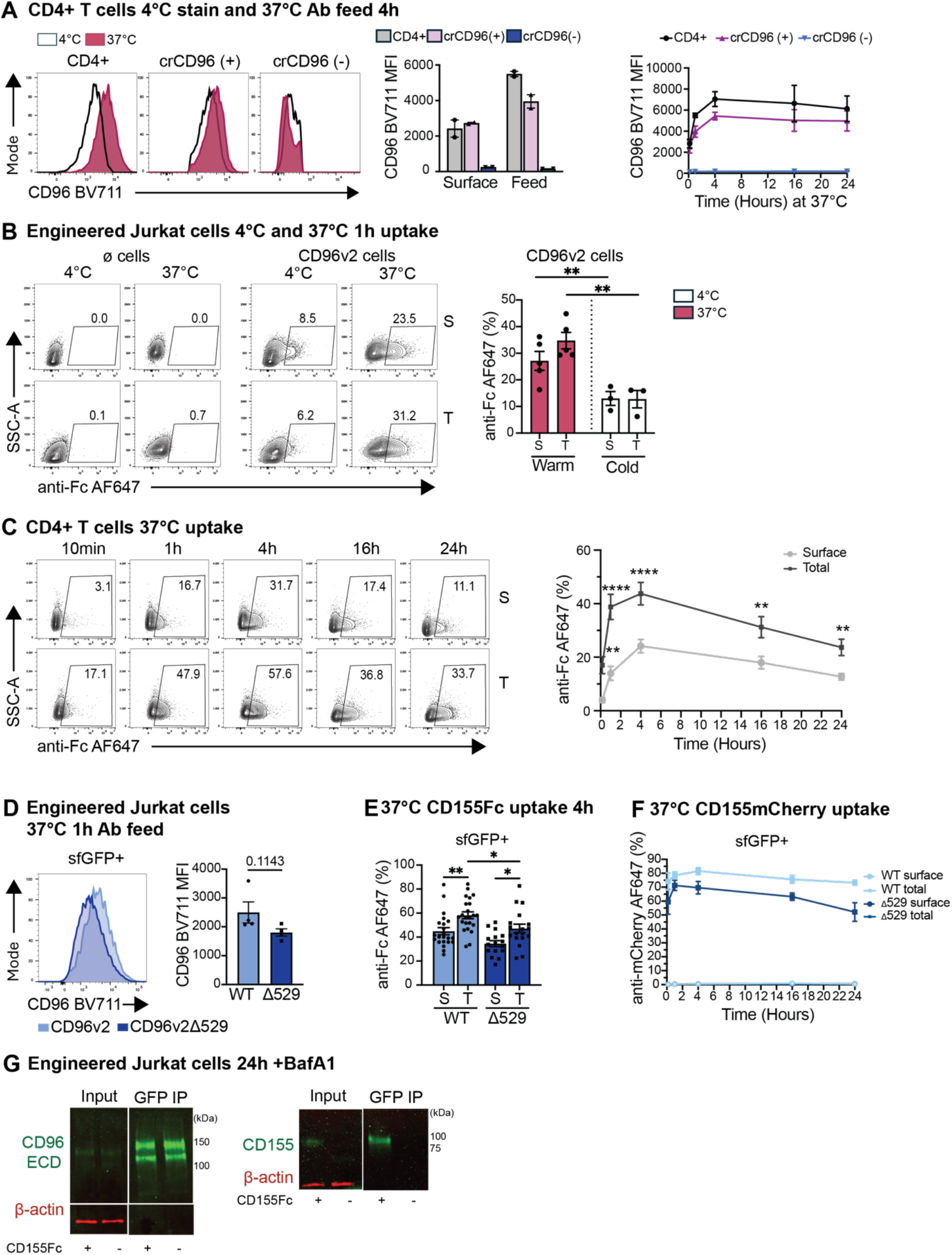
CD96 cycles and internalises CD155 in an active process. **(A)** CD4+ or CRISPR-Cas9-mediated knockout of CD96 (crCD96), expanded, primary human CD4+ T cells were incubated with anti-CD96 BV711 antibody for antibody feed for times as specified at 37°C and 5% CO_2_ or for surface stain for 4h at 4°C (n=2 each, matched). Representative histograms (left) and median fluorescence intensity (MFI, middle) at 4h, and time-course (10min, 1h, 4h, 16h, and 24h at 37°C and 5% CO_2_, right) for anti-CD96 BV711 MFI (CD4+ CD96+/-gated). **(B)** Engineered Jurkat cells without (ø)/with CD96v2sfGFP expression were incubated with soluble CD155Fc (5μg/mL) for 1h at 37°C and 5% CO_2_ or at 4°C, stained with anti-human Fc AF647 antibody on the surface or post-fixation/permeabilisation for the total stain. Representative plots and frequency of the surface and total anti-human Fc AF647 (CD155) (n=5). Data as mean±SEM and two-way ANOVA with Bonferroni multiple comparisons. **(C)** Expanded primary human CD4+ T cells were incubated with soluble CD155Fc (5μg/mL) for 10min, 1h, 4h, 16h, and 24h at 37°C and 5% CO_2_, stained with anti-human Fc AF647 antibody on the surface or post-fixation/permeabilisation for the total stain. Representative plots and frequency of the surface and total anti-human Fc AF647 (CD155) (n=9). Data analysed using paired (Wilcoxon) test for each timepoint. **(D)** Engineered Jurkat cells expressing WT CD96v2sfGFP or CD96v2 with cytoplasmic domain deletion (CD96v2Δ529sfGFP) were incubated with anti-CD96 BV711 for antibody feeding for 1h at 37°C and 5% CO_2._ Representative histogram (sfGFP+ gated) and MFI of anti-CD96 BV711 staining (n=4). Data as mean±SEM and Mann-Whitney test. **(E-F)** Engineered Jurkat cells expressing WT CD96v2sfGFP or CD96v2Δ529sfGFP were incubated with soluble CD155Fc (5μg/mL, shown at 4h with one-way ANOVA with Bonferroni multiple comparisons, **E**) or 1:1 with DG75.CD155mCherry (**F**) for 10min, 1h, 4h, 16h, and 24h at 37°C and 5% CO_2_, stained with anti-human Fc or anti-mCherry AF647 antibody on the surface or post-fixation/permeabilisation for the total stain. Frequency of surface and total CD155Fc (**E**, n=24) or CD155mCherry (**F**, n=9) stained on sfGFP+ gated cells. Data as mean±SEM. **(G)** Engineered Jurkat cells expressing CD96v2sfGFP were incubated with soluble CD155Fc (5μg/mL) ± Bafilomycin A1 (BafA1, 30nM) for 24h at 37°C and 5% CO_2._ Cells were lysed, GFPtrap (GFP IP) and input were assessed for CD96 (extracellular domain, ECD), CD155, and house-keeping (beta-actin) using WB. Precision plus protein ladder used, molecular mass (kDa) shown. **p<0.01, ****p<0.0001.

We confirmed active cycling using our engineered Jurkat cells expressing CD96v2 incubated with CD155Fc which showed greater surface and total CD155Fc at 37°C compared to 4°C (p=0.0012, **Figure 4B**). Notably, total CD155Fc was greater than surface at 37°C (34.72±3.08 vs 27.12±2.56) but not at 4°C (12.75±3.25 vs 12.99±2.61), confirming that CD96-dependent internalisation of CD155 is an active process. Time-dependent internalisation of CD155 was also observed in CD4+ T cells, with surface and total CD155Fc peaking at 4h (**Figure 4C**), mirroring anti-CD96 antibody feed kinetics (**Figure 4A**). Unlike anti-CD96, which was present in excess, CD155Fc uptake decreased after 4h, suggestive of ligand degradation when ligand is limited.

To assess the role of cytoplasmic region in CD96 cycling, GFP-matched CD96v2sfGFP expressing cells were compared to cells with a cytoplasmic domain deletion (CD96v2Δ529, **supplemental Figure S2**). WT CD96v2 cells exhibited higher anti-CD96 BV711 MFI in an antibody feed than CD96v2Δ529 cells (MFI: 2497.0±365.8 and 1803.0±124.9 respectively, **Figure 4D**), demonstrating that the cytoplasmic domain contributes to receptor cycling. Indeed, CD155Fc (**Figure 4E, supplemental Figure S3A**) and cell-expressed CD155mCherry (**Figure 4F, supplemental Figure S3B**) uptake assays showed reduced internalisation with cytoplasmic domain deletion. At 4h, WT CD96v2 expressing cells displayed greater total CD155Fc staining compared to CD96v2Δ529 (60.4%µ4.8 vs 41.0%µ2.7, p=0.0466, **Figure 4E**), indicating impaired ligand uptake when the cytoplasmic domain is deleted. Interestingly, in cell-cell transfer of CD155mCherry, no surface anti-mCherry was observed (**Figure 4F**), demonstrating that CD155 is actively transferred across membranes at synapses.

Probing a GFPtrap-IP (CD96v2sfGFP Jurkat culture ± CD155Fc) for CD155 confirmed direct CD96-CD155 interaction (**Figure 4G**). The CD96 (left, **Figure 4G**) and CD155 (right, **Figure 4G**) WB bands demonstrate successful CD96 pulldown and co-IP of CD155Fc and reinforce that active binding and internalisation of CD155 is CD96-dependent. Collectively, these results demonstrate that CD96 actively cycles, partially facilitated via its cytoplasmic domain, and mediates internalisation of both soluble and cell-expressed CD155 through direct interaction.

### Internalised CD155 undergoes lysosomal degradation

We have shown here that internalised CD155 decreased over time (see **Figure 4**), thus suggesting that the ligand undergoes degradation upon internalisation. CD4+ T cells treated with degradation inhibitor Bafilomycin A1 (BafA1) showed an accumulation of CD155 at the later timepoints, (total CD155Fc stain at 16h 55.80±2.72% vs 32.98±4.07%, p=0.0006; surface: 32.45±1.88% vs 18.72±2.45%, p=0.0001, **Figure 5A**) and at 24h (total: 49.14±4.54% vs 23.13±2.28%, p=0.0002; surface: 27.88±3.37% vs 12.60±0.92%, p=0.0082, **Figure 5A**). Similarly, confocal microscopy imaging showed accumulation of internalised CD155Fc in engineered CD96v2sfGFP expressing HeLa cells treated with degradation inhibitor ammonium chloride (NH_4_Cl) (number of CD155Fc puncta per cell 68.38±4.71 vs 56.66±4.50, p= 0.0002, **Figure 5B**). Puncta also appeared larger and more prominent (**Figure 5B**) indicating preservation of internalised ligand in degradation vesicles.

**Figure 5.**
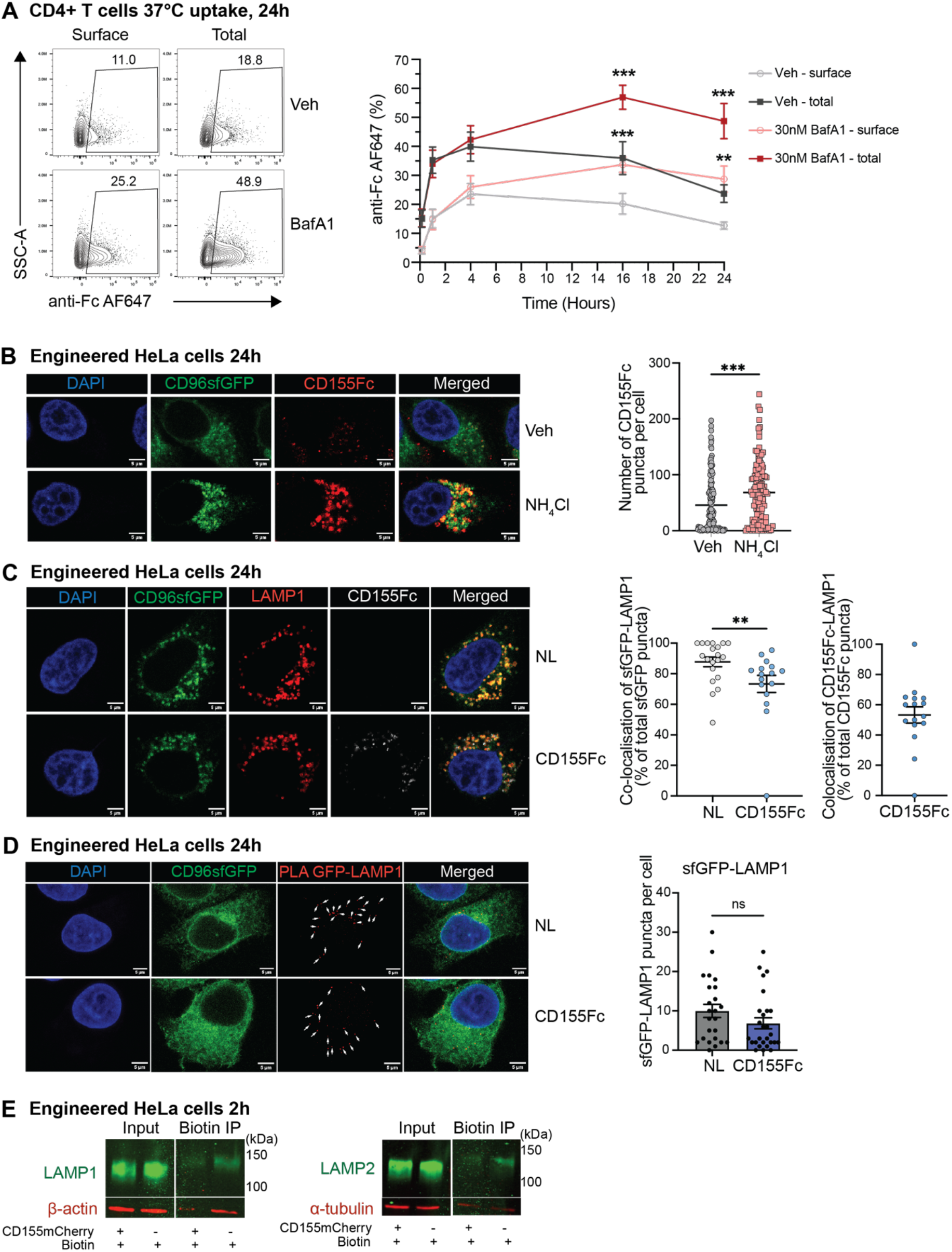
Internalised CD155 undergoes lysosomal degradation. **(A)** Expanded primary human CD4+ T cells were incubated with soluble CD155Fc (5μg/mL) ± Bafilomycin A1 (BafA1, 30nM) or vehicle (Veh) for 10min, 1h, 4h, 16h, and 24h at 37°C and 5% CO_2_, stained with anti-human Fc AF647 antibody on the surface or post-fixation/permeabilisation for the total stain. Representative plots (of 24h) and frequency of the surface and total anti-human Fc AF647 (CD155) (n=6). Data as mean±SEM and one-way ANOVA with Holm-Šídák’s multiple comparisons test between Veh vs BafA1 for each timepoint. **(B)** Engineered HeLa cells expressing CD96sfGFP (green) incubated with soluble CD155Fc (5μg/mL, anti-human Fc AF647 antibody, red) ± NH_4_Cl (30mM) for 24h at 37°C and 5% CO_2_; with DAPI (blue) stain. Representative confocal microscopy images and quantification of number of CD155Fc puncta (red) per cell. Quantification of 53 images in total from two independent experiments. **(C)** Engineered HeLa cells expressing CD96sfGFP incubated ± soluble CD155Fc (5μg/mL) for 24h at 37°C and 5% CO_2_, with DAPI (blue) stain. Representative confocal microscopy images and frequency of co-localisation between CD96sfGFP (green)-LAMP1 (red) (of total sfGFP puncta) and CD155Fc (grey)-LAMP1 (of total CD155Fc puncta, far right). Analysis of 3-4 images per condition from 2 experiments. **(D)** Engineered HeLa cells expressing CD96sfGFP (green) incubated ± soluble CD155Fc (5μg/mL) for 24h with DAPI (blue) stain and proximity ligation assay (PLA) assessing direct interaction (<40nm, seen as red puncta) between sfGFP and LAMP1 with quantification of number of puncta per cell. Image quantification for each cell from 3 images per condition taken from one experiment. **(E)** Engineered HeLa cells expressing CD96miniTurboID were incubated 5:1 ± CellTrace blue labelled (CTB) DG75.CD155mCherry cells and with biotin (300uM) for proximity labelling (<10nm) for 2h. Cells were lysed and streptavidin-agarose biotin IP and input assessed for LAMP1 or LAMP2 (green), and housekeeping (beta-actin/alpha-tubulin; red) using WB. Precision plus protein ladder used, molecular mass (kDa) shown. Scale bar throughout=5μm; Data as mean±SEM; **p<0.01, ***p<0.001.

To ascertain the mode of degradation, co-localisation between CD96sfGFP and the lysosomal protein LAMP1 was assessed, with significant co-localisation of LAMP1 with CD96sfGFP (87.74±3.14% of all CD96sfGFP puncta, **Figure 5C**) and CD155Fc (53.24±5.37% of all CD155Fc puncta, **Figure 5C**), indicating lysosomal degradation of co-receptor and ligand. Proximity ligation assay (PLA, measuring protein-protein interaction within 40nm) confirmed CD96-LAMP1 interaction (12.56±2.89-15.58±2.48 PLA puncta, **Figure 5D**). Of note, in both confocal microscopy co-localisation and PLA CD96 co-localised with LAMP1 regardless of ligand presence (87.74±3.14% vs 73.32±5.58%, p=0.0064, **Figure 5C;** 10.00±1.67 vs 6.85±1.29, p=0.1475, **Figure 5D**) suggesting a constant lysosomal degradation pathway for CD96. This was further confirmed by miniTurboID proximity labelling (capturing interactions within 10nm at CD96 C-terminus) for IP-WB, which validated lysosomal LAMP1 and LAMP2 as CD96 interacting partners (regardless of ligand presence, **Figure 5E**).

Collectively, these findings demonstrate that CD96 constantly cycles and traffics to lysosomes to be degraded and directs CD155 into the same degradative pathway when bound and internalised.

### CD155 may enter selective autophagy pathways via CD96 interaction

Cell cargo can arrive at lysosomes directly via endosomes or directed through autophagosomes^28, 29^. CD155Fc co-localised with the key autophagy protein LC3 in engineered CD96v2sfGFP expressing HeLa cells by confocal microscopy, which significantly increased in the presence of the degradation inhibitor NH_4_Cl (19.23±3.86% vs 36.77±3.16%, p=0.0014, **Figure 6A**). Additionally, CD155Fc co-localised with the selective autophagy protein p62, which also significantly increased following NH_4_Cl treatment (36.44±4.90% vs 61.92±2.72%, p=0.0001, **Figure 6B**). These findings indicate that CD155Fc accumulates in autophagosomes/autophagolysosomes when degradation is inhibited. Interestingly, CD96v2sfGFP showed strong co-localisation with LC3 and p62, which was not affected by the NH_4_Cl treatment (LC3 co-localisation: 51.33±7.49% v 41.41±3.98%, p=0.3184, **Figure 6A**; p62 co-localisation: 63.77±5.71% vs 76.19±3.45%, p=0.1829, **Figure 6B**). CD96-LC3 and CD96-p62 interactions were also confirmed by PLA (**Figure 6C**). Whilst these data indicate CD96 trafficking into degradation may be facilitated by selective autophagy, miniTurboID-IP-WB did not detect LC3 or p62 as direct CD96 interactors (**Figure 6D**), indicating a possible unidentified bridging protein linking CD96 to LC3 and p62. However, autophagy related proteins ATG5 and ATG7 were detected (**Figure 6E**), indicating close interaction and entry of CD96 into the autophagy pathway. Thus, internalised CD96 and CD155 can enter selective autophagy for degradation in autolysosomes.

**Figure 6.**
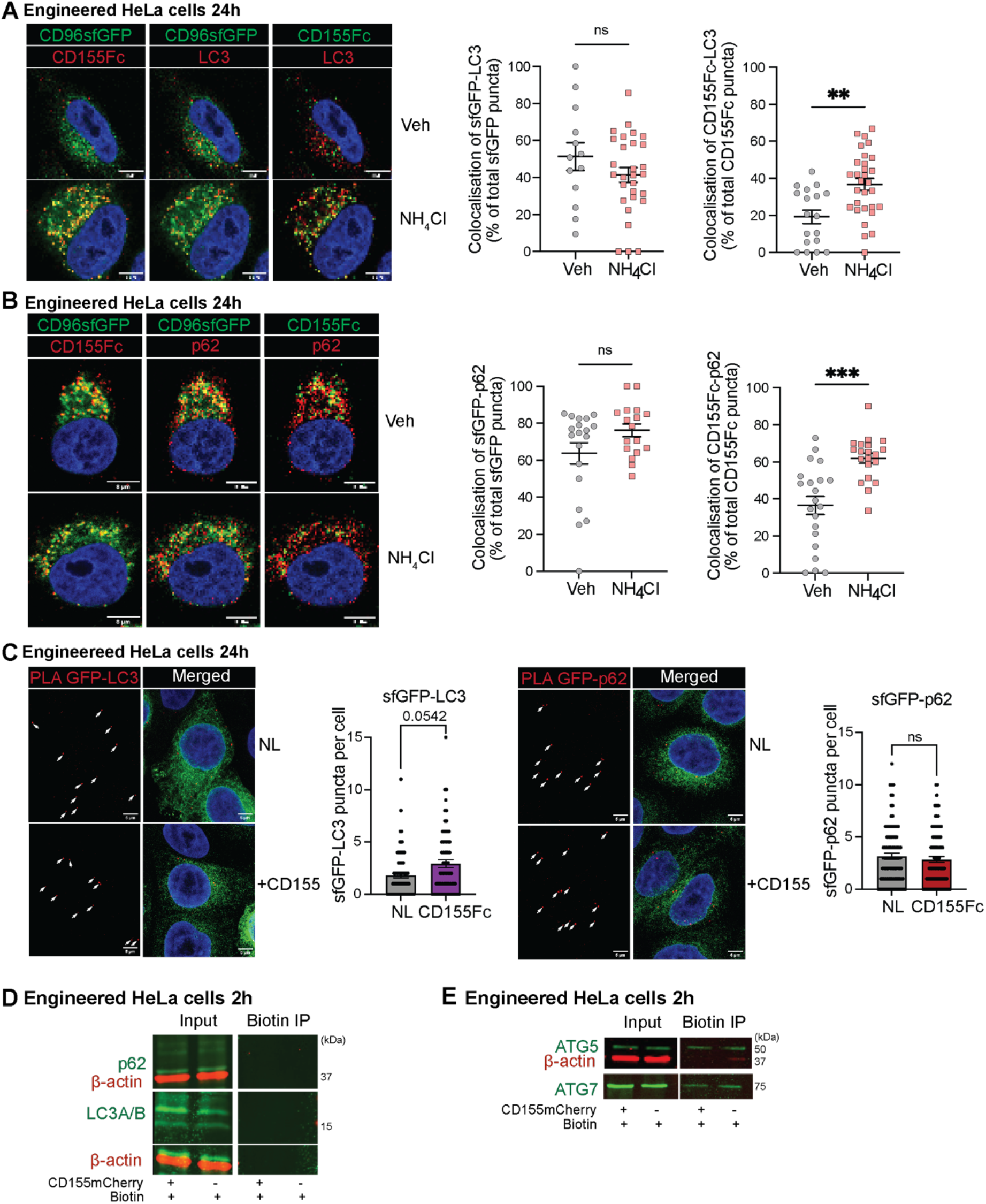
CD96 and CD155 interaction with selective autophagy pathway markers. **(A-C)** Engineered HeLa cells expressing CD96sfGFP were incubated ± soluble CD155Fc (5μg/mL) for 2h or 24h ±NH_4_Cl or vehicle (Veh) at 37°C and 5% CO_2_. **(A)** Confocal microscopy representative images and frequency of CD96sfGFP (green)-LC3 (red) (of total sfGFP puncta) and CD155Fc (green)-LC3 (red) (of total CD155Fc puncta). **(B)** Confocal microscopy representative images and frequency of CD96sfGFP (green)-p62 (red) (of total sfGFP puncta) and CD155Fc (green)-p62 (red) (of total CD155Fc puncta). **(C)** Proximity ligation assay (PLA) assessing direct interaction (<40nm, seen as red puncta) between sfGFP and LC3 or p62 with the number of puncta per cell in each condition (NL=no ligand). **(D-E)** Engineered HeLa cells expressing CD96miniTurboID were incubated 5:1 ± CellTrace blue labelled DG75.CD155mCherry cells and with biotin (300uM) for proximity labelling (<10nm) for 2h. Cells were lysed and streptavidin-agarose biotin IP and input assessed for p62 and LC3A/B (**D**; green), or ATG5 and ATG7 (**E**; green) and house-keeping (beta-actin; red) using WB. Precision plus protein ladder used, molecular mass (kDa) shown. Scale bar throughout=8μm; Data as mean±SEM and Mann-Whitney test; **p<0.01, ***p<0.001.

## Discussion

CD96 is a member of the immunoglobulin superfamily consisting of TIGIT, CD226, CD96 co-receptors and their shared ligand, CD155, that has been implicated in immune regulation^1, 3, 4^. The functional role of CD96 remains incompletely defined compared with the more characterised co-inhibitory and co-stimulatory functions of TIGIT and CD226, respectively. Conflicting reports on CD96 function across immune and disease contexts suggest that its activity may depend on its interaction with shared ligands and co-receptor expression within the local microenvironment.

In this study, we investigated CD96-CD155 interactions and reveal a mechanism by which CD96 regulates ligand availability through receptor-mediated internalisation and degradation. We identified CD96 as a broadly expressed co-receptor across T cell subsets and demonstrate that it uniquely facilitates the uptake and internalisation of its shared ligand CD155 within the TIGIT/CD226/CD96 co-receptor family. Our data indicate that this is dependent on active receptor cycling and directs the CD155 towards lysosomal degradation pathways, with evidence supporting involvement of selective autophagy. These findings reveal a previously unrecognised mechanism by which CD96 may regulate ligand availability, with implications for T cell function and immune regulation.

The emerging upregulation of CD96 and its ligand CD155 across inflammatory and malignant settings highlight their potential clinical relevance. Increased CD96 expression is observed in various disease settings such as autoimmune vasculitis, inflamed joints in juvenile idiopathic arthritis, and multiple malignancies where it is associated with poor prognosis^5–7^. Similarly, CD155 is frequently upregulated in several malignancies including colorectal cancer and lung adenocarcinoma and is associated with adverse outcomes^2, 8, 9^. Taken together, these observations highlight the CD96-CD155 complex, as well its associated receptors TIGIT and CD226, as a compelling target for therapeutic intervention, particularly in cancer immunotherapy whereby modulation of immune checkpoint pathways has shown significant clinical benefits^3, 30^.

We establish a novel mechanism by which CD96 mediates the internalisation and degradation of the ligand and thus build on limited previous literature which has suggested possible transfer of CD155 onto CD96+ cells. Fuchs et al showed example flow cytometry histograms and immunofluorescence images of CD155 on CD96+ NK92 cells after co-culture with CD155 transfected cells^31^. Similarly, Huang et al also showed some evidence for CD155 transfer to T cells which was reduced upon CD96 blockade or knockdown^32^. However, neither of these studies investigated underlying mechanisms.

We demonstrate that CD96+ cells are capable of uptake and internalisation of both cell-bound and soluble CD155. This is particularly of importance as CD155 exists as different isoforms, comprising two membrane-bound forms (α and δ) and two soluble forms (β and γ)^2^, all of which CD96 would be capable of regulating. Soluble CD155 has also been reported to be elevated in serum of various malignancies and is associated with tumour burden and advanced cancer stages^33, 34^. Furthermore, we demonstrated CD96-mediated uptake of CD155 across multiple CD96-expressing cell types, including primary CD4+ T cells (Tconvs, Tregs) and model cell lines (engineered Jurkat and HeLa cells), suggesting that this function of CD96 is conserved across different cell types.

Mechanistically, the data suggests that CD96 undergoes constitutive trafficking to lysosomal and autophagy pathways. Antibody feeding experiments together with the interaction with lysosomal proteins (LAMP1, LAMP2) and autophagy-related proteins (ATG5, ATG7) suggests that CD96 is undergoing continuous active cycling through degradative compartments, during which CD155 is internalised. The complex undergoes degradation via the selective autophagy pathway as seen through interaction between CD96, CD155, and autophagy proteins ATG5, ATG7, LC3 and p62. Although we did not observe direct interaction (1-10nm) between CD96 and selective autophagy proteins p62 or LC3, we revealed interactions within 40nm, suggesting an indirect interaction with CD96, such as an unknown mediator protein in-between. Our findings are consistent with previous work by Huttlin et al, who, using high-throughput affinity purification–mass spectrometry, identified interaction between CD96 with CD155 and autophagy-associated proteins such as p62, as well as endocytosis-associated Hsp70 and E3 ubiquitin ligase proteins KCMF1 and URB4, with these CD96-related interactions reported within the additional dataset of their global interactome analysis^35–37^. Interestingly, we also observed that CD155 uptake was impaired by CD96 cytoplasmic domain deletion. Some internalisation was maintained with truncated CD96, potentially due to high ligand density and crowding in co-culture from highly overexpressed CD155 and CD96 on modified cell lines. Nonetheless, the impaired CD155 uptake by truncated CD96 indicates the importance of CD96 residues with the cytoplasmic domain in facilitating ligand uptake. Receptor-mediated endocytosis can be dynamin-dependent such as clathrin-mediated e.g. for CTLA-4^38^ whereby adaptor protein 2 complex (AP2) facilitates specific cargo binding^39, 40^, or clathrin-independent^41^. Notably, CD96 cytoplasmic domain contains two YXXΦ motifs (YHEM, YTCI), which are putative AP2-binding sites that could mediate clathrin-dependent endocytosis^39, 42^. The motifs may also interact with AP1 or AP3 complexes, implicated in endosomal trafficking and degradation respectively^43, 44^.

Taken together our data provides clear evidence for the hypothesis that CD96 regulates the immune response by regulation of ligand availability rather than via mechanisms such as classical downstream signalling. Modulating the availability of CD155 for TIGIT and CD226 will alter the immune response, since TIGIT acts as co-inhibitory and CD226 as co-stimulatory co-receptor^3^, thus CD96 indirectly affects immune cell activation and function. This mechanism also helps explain the discrepancies in previous literature regarding the role of CD96 as a co-inhibitory or co-stimulatory receptor^7, 15–17, 19 18^, which may depend, at least in part, on expression of TIGIT and CD226 within the local microenvironment. For instance, the reduction in tumour burden observed with anti-CD96 treatment by CD8+ T cells, as reported by Mittal et al.^17^, may also be explained by ligand availability. Blocking the CD96-CD155 interaction with an anti-CD96 antibody would increase the availability of CD155 for the co-stimulatory receptor CD226, thereby enhancing CD8+ T cell activation and promoting anti-tumour responses. Consistent with this interpretation, Mittal et al. showed that the anti-tumour effect of CD96 blockade was reversed when CD226 was blocked or absent (in CD226^−^/^−^ mice)^17^, supporting the model that the observed effect is partly mediated by increased ligand availability for CD226. Further competition-based experiments will be required to directly assess the impact of CD96 mediated ligand regulation on the functional output of cells expressing TIGIT or CD226.

In summary, this study reveals a previously unrecognised role for CD96 in modulating CD155 availability through constitutive receptor cycling and lysosomal/autophagy-associated degradation. By modulating ligand access to TIGIT and CD226, CD96 may act as a key determinant of immune balance within this receptor axis. These findings provide a mechanistic basis for the context-dependent roles of CD96 described in the literature and may inform future therapeutic targeting strategies.

## Methods

### Study approval

This research was conducted with ethical approval (Royal Free London NHS Foundation Trust (RFL)-UCL biobank REC Reference Number: 21/WA/0388, IRAS Project ID: 309315, RFL Biobank Ethical Reference numbers: NC2023.17/NC2023.20; UCL research ethics 14017/001, 14017/002) and informed consent in accordance with the Declaration of Helsinki.

### Tissue culture, cell lines and cell line engineering

Primary cells were cultured in complete Optimizer media (CTS™ OpTmizer™ + 2.54% supplement, 100U/mL penicillin-streptomycin, and 1% GlutaMAX (all ThermoFisher)), unless indicated otherwise in the text.

All cell lines were grown and maintained at 37°C and 5% CO_2_. Jurkat, DG75 and L cell lines were maintained in complete RPMI1640 media (ThermoFisher) supplemented with 10% foetal bovine serum (FBS; Life Science Production), 100 U/mL penicillin-streptomycin, and 1% GlutaMAX (all ThermoFisher). HeLa cell lines were maintained in complete DMEM media + 10% FBS, 1% GlutaMAX, and 1% penicillin-streptomycin.

DG75.2KO (previously KO for CD80 and CD86 using CRISPR-Cas9, provided by Sansom lab), Jurkat, and HeLa cell lines were CRISPRed (as below) for co-receptors (TIGIT/CD226/CD96) and ligands (CD155/CD112), pure populations were sorted, and further engineered by lentiviral-mediated overexpression of molecules of interest as indicated in the text and populations purity sorted. The phenotype of engineered cell lines was regularly confirmed by flow cytometry.

Lentiviral particles were produced by transfection of Hek293T cells with transfer plasmid, VSV-G, and 8.91 packaging by calcium phosphate^45^. Media was exchanged at 12-16h and supernatant was collected after 48h further culture, titre assessed on Hek293T and frozen at −70°C in aliquots.

Transfer plasmids were based on pDual_GFP (Addgene 86980, provided by the Grove lab), which was modified to replace BAMHI cloning side with BstZ17I after the SFFV promoter and removing GFP reporter sequence (resulting in pDualB, Pesenacker lab). CD96 and CD155 codon optimised open reading frames were purchased (ThermoFisher) and cloned into pDualB with STOP codon or with flexible linker^46^ fused to tags (sfGFP/mCherry/CFP/miniTurboID) using standard restriction enzyme digestion-ligation (NEB reagents). Variants were cloned using standard PCR-cloning, Q5-mediated site directed mutagenesis (NEB) or restriction enzyme digestion-ligation.

### Blood processing, T cell isolation, and expansion

Peripheral blood (PB) or non-clinical-issue leukocyte cones (LC, NHSBT) were processed to PBMCs or CD4+ T cells by pre-incubating PB/LC with RosetteSep™ Human CD4+ T Cell Enrichment Cocktail (12.5µL/mL PB, 5µL/mL LC, Stemcell technologies) via density gradient centrifugation according to standard protocols and cryopreserved.

CD4+ T cells were enriched/depleted for CD25 using CD25 MicroBeads-II (Miltenyi Biotec) and the positive fraction purity sorted for Tregs (live (FVD-) CD4+CD25hiCD127lo) and the negative fraction purity sorted for Tconvs (live (FVD-) CD4+CD25loCD127hi) by flow cytometry on BD FACSAria Fusion.

T cells were expanded on L cells (artificial antigen presenting cells (aAPC), provided by Levings Lab) mouse fibroblasts overexpressing human CD80, CD32, and CD58^47^; loaded with 100ng/mL mouse anti-human CD3 (clone OKT3; Biolegend) in complete OpTmizer with 100IU/mL (for CD4+ T cells and Tconvs) or 1000IU/mL (for Tregs) IL-2 (Proleukin) at 37°C and 5% CO_2_ for 5-7 days.

### CRISPR-Cas9 mediated knockout (KO)

Primary T cells were pre-activated using ¼ of the recommended CD3/CD28/CD2 tetramer (Stemcell) for 2 days before a CRISPR-Cas9 mediated KO. Briefly, guide RNAs (crRNA:tracrRNA duplex, IDT) were designed for each target and complexed with recombinant Cas9 protein (provided by Marson lab,UCSF or IDT) for 10-20min at RT and combined with cells in the buffer T/R (primary cells/cell lines), and electroporation using the Neon^TM^ Transfection System (ThermoFisher) with optimised settings (primary T cells: 2000V, 20 width, 2 pulse; Jurkat: 1700V, 20 width, 1 pulse; DG75: 1600V, 10 width, 3 pulse; Hela: 1300V, 20 width, 2 pulse). Cell lines were cultured as described above and primary T cells were expanded on L cells for 5-7 days as described above before CRISPR efficiency was assessed/purity sorted by flow cytometry.

### In vitro assays

*Antibody feed:* Cells were incubated with anti-CD96 for 10min, 1h, 4h, 16h, and 24h at 37°C and 5% CO_2,_ before being processed for flow cytometry.

*CD155 uptake assays:* For soluble CD155 uptake assays, primary T cells or modified Jurkat cell lines were incubated with 5ug/mL recombinant human CD155/PVR protein with human Fc tag (CD155Fc; SinoBiological) for 10min, 1h, 4h, 16h, and 24h at 37°C and 5% CO_2_, or incubated with CD155Fc for 1h at 4°C. Cells were stained with anti-human Fc monoclonal antibody and relative antibodies/dyes (*supplemental Table S1*) for flow cytometry, confocal microscopy or IP-WB.

For uptake assays from ligand expressing cells, CD96 expressing engineered HeLa or Jurkat cells with or without C-terminal tags were co-cultured at different ratios as specified in legends with CellTrace Blue labelled engineered DG75 cells expressing CD155mCherry or CD155CFP. Cells were stained with anti-mCherry and relative antibodies/dyes (*supplemental Table S1*) for flow cytometry, confocal microscopy or IP-WB. Where indicated, Bafilomycin A1 (BafA1, 20-30mM; Stratech) or Ammonium Chloride (NH_4_Cl, 10-30mM; Fisher) degradation inhibitors were used. Biotin (0 or 300uM; Cambridge Biosciences) was added for miniTurboID proximity labelling for last 2h. Where indicated, DSP cross-linker (dithiobis(succinimidyl propionate), 5mM; Fluorochem Ltd) was added for 30min at room temperature and quenched using Tris-HCl (25mM; ThermoFisher) for 15min.

### Flow cytometry

Cells were stained on the surface with antibodies (see *supplemental Table S1*) and viability dye for 30min at 4°C. For CD155 uptake assays, surface staining included anti-human Fc or anti-mCherry antibody to detect ligand on the surface. Cells were fixed and permeabilised with Foxp3 Transcription Factor Fixation/Permeabilization buffer (eBioscience^TM^) as per the manufacturer’s instructions and split before half of the samples were stained with intracellular antibodies for 45min at 4°C, including the addition of anti-human Fc and anti-mCherry antibodies to assess the total surface and intracellular level of ligand. Samples were acquired using BD LSRFortessa^TM^ X-20 or Cytek Aurora. Antibodies and dyes used for flow cytometry are listed in *supplemental Table S1*.

### CD96 gene expression

mRNA expression of CD96 variants was performed by nanoString. Cell lysates (RNeasy Lysis Buffer (RLT; Quiagen) with 1% 2-Mercaptoethanol at 1μL/5000 cells) were hybridised to the Pesenacker Hu_TregsPlus custom nCounter CodeSet and analysed using the nanoString nCounter. QC and normalisation included negative control check, positive control normalisation factor and total sum normalisation factor followed by log2(1+x) transformation^48^.

### Immunoprecipitation (IP) and western blot (WB)

Uptake assays were carried out as indicated, cells washed with PBS and lysed with fresh ice-cold RIPA lysis buffer for 30min with rotation at 4°C. The soluble fraction was carried into immunoprecipitation (IP) or stored at −70°C for SDS-page-WB.

Streptavidin-agarose beads (Merck) and Chromotek GFPtrap beads (Proteintech) were used according to manufactures guidelines. IP beads were incubated with fresh cell lysates (5M cells) for 2h with rotation at 4°C, washed 3x before elution in 40-100uL 4xLaemmli loading buffer (BioRad) with reducing agent DTT (50mM, dithiothreitol; ThermoFisher) and boiling for 5-10min at 95°C. Sample were stored at −70°C and beads sedimented using centrifugation before SDS-page-WB.

Samples (lysate input, IP) were loaded into 4-20% Mini-PROTEAN TGX precast protein gels (BioRad) and run for 50min at 140V before semi-dry transfer (Trans-Blot, BioRad) onto activated Immobilin-FL PVDF membranes (Merck) for 7min at 25V, 2.5A. Membranes were air-dried, then blocked using Intercept Blocking Buffer (LI-COR) for 1h at RT, and stained in staining buffer overnight (16-18h) at 4°C. After 5x washes, membranes were stained with secondary antibody for 1h at RT and washed 5x before visualisation using Odyssey M imager (LI-COR). Antibodies used for WB are listed in *supplemental Table S1* and buffer recipes *supplemental Table S2*.

### Confocal microscopy

Engineered cells were labelled (where indicated) and co-cultured as indicated (recombinant ligand or ligand-expressing cells) on imaging compatible 96-well plates (Griener), for suspension cells poly-L-lysine–coated, and for Jurkat cells in the presence of 1µg/mL αCD3 (OKT3; Biolegend), for indicated time between 2-24h at 37°C. For inhibition of lysosomal acidification, bafilomycin A1 (20nM) or ammonium chloride (30mM) was added as indicated. Cells were fixed by centrifugation with 2% paraformaldehyde (PFA; Fisher) for 15-20min at RT. For antibody labelling, cells were permeabilised and blocked in PBS containing 5% bovine serum albumin (BSA; Merck) and 0.1% Triton™ X-100 (Merck) for 1h at RT. Cells were incubated with primary antibodies overnight at 4°C (see *supplemental Table S1*). Cells were stained with secondary antibodies for 1-2h at 4°C. Where indicated, cells were stained with DAPI (1 μg/mL; Merck) for 5min at RT. Cells were washed, mounted in Mowiol mounting medium containing 2.5% 1,4-diazobicyclo-[2.2.2]-octane (DABCO, Sigma), and stored at 4°C protected from light or imaged immediately by confocal microscopy on Nikon Ti Eclipse C2 laser scanning confocal microscope using 60x oil immersion objective.

*Proximity ligation assay (PLA):* Engineered HeLa cells were incubated as above for 24h before fixation with 4% PFA for 15min at RT. Cells were washed and permeabilised (PBS with 5% BSA, 0.1% Triton X-100) for 1h at RT. Proximity Ligation Assay (PLA) was performed using the Duolink® In Situ Red Starter Kit (Sigma-Aldrich) according to the manufacturer’s instructions. Briefly, samples were blocked with Duolink® Blocking Solution for 1h at 37°C and incubated overnight at 4°C with the relevant primary antibodies (see *supplemental Table S1*) in Duolink® Antibody Diluent. With appropriate washing (Wash Buffer A) between steps, PLUS and MINUS PLA probes (anti-mouse and anti-rabbit) were applied for 1h at 37°C. Ligation was performed using ligation buffer containing ligase (1:80 dilution) for 30min at 37°C. Amplification buffer containing polymerase (1:160 dilution) was added for 100min at 37°C protected from light. Then, samples were washed twice in Wash Buffer B. Minimal volume (1-2 drops) of Duolink® In Situ Mounting Medium containing DAPI was added and imaged by confocal microscopy (Nikon Ti Eclipse C2) after 15min. Antibodies for confocal microscopy are listed in *supplemental Table S1*.

### Analysis

Flow cytometry data was analysed using FlowJo v10 (BD) for manual gating (cells, single cells, FVD-live cells and as indicated) and marker expression assessment.

Publicly available spectral flow cytometry data (FLOWRepository ID FR-FCM-Z6VC)^6^ was analysed using unbiased clustering analysis via R with R package Spectre^26^ using PhenoGraph clustering algorithm^25^.

Confocal images were visualised using FIJI (ImageJ) and raw images were analysed and quantified using CellProfiler^TM^. Images were segmented using DAPI and GFP to identify the cells. GFP, CD155Fc, and LC3/p62/LAMP1 markers were also segmented as secondary objects or puncta. The frequency of co-localisation was calculated as the percentage of overlap between the LC3/p62/LAMP1 puncta over total sfGFP or CD155Fc puncta per cell as previously described^49^. For PLA analysis, the red PLA puncta were identified and counted per cell.

Statistical analyses were performed using Graphpad Prism v10.4.1: unpaired (non-parametric Mann-Whitney) or paired (Wilcoxon) test for comparisons of two groups, one-way ANOVA with Bonferroni or Holm-Šídák’s multiple comparisons test for >3 groups, and two-way ANOVA with Bonferroni multiple comparisons or Šídák’s multiple comparisons test for multiple grouped comparisons. Correlation analysis was performed using nonparametric Spearman correlation and a simple linear regression. Welch’s t test was used to compare AUCs. p values <0.05 were considered statistically significant, and data presented as mean ± standard error of the mean (SEM) throughout.

## Acknowledgments

The authors would like to thank the Levings Lab (UBC BC-CHRI), Sansom Lab (UCL IIT), Grove Lab (University of Glasgow), Marson Lab (UCSF), and Pallett Lab (UCL IIT) for contributing to this research by providing us with various cell lines and reagents. We would like to thank all the healthy donors for volunteering blood samples for our research. We would like to thank Dr Claudia Hinze for her support with confocal microscopy analysis. We acknowledge and thank Janani Sivakumaran Nguyen and Sam Blanchett at the IIT flow cytometry facility for all their technical help and support. We would like to thank Prof David Sansom and Prof Lucy Walker and the teams at UCL IIT for all the discussion, feedback, and exchange of ideas.

## Author Contribution

DS and RF contributed to overall study design, execution of all experiments, data analysis, writing and editing of the manuscript. MK, RW contributed to execution of experiments and data analysis. MHA, RS assisted with execution of experiments, data analysis, and editing of the manuscript. AMP led and oversaw the overall study design, data acquisition, analysis and writing and editing of the manuscript, and takes responsibility for the content of the article. All authors read and approved the manuscript.

## Funding

This project was funded by UKRI BBSRC BB/V009524/1. Arthritis UK (CDF21738, 23159, 23135), NIHR BRC UCLH (BRC764/III/AP/101350) and Kidney Research UK (SG_MNRP_006_20241015) supported salaries of AMP, DS, MHA, and RS, and contributed to general shared lab consumables. MRC DTP PhD studentship (MR/W006774/1) supported the stipend for RF and contributed to general shared lab consumables.

## Notes

### Competing Interest Statement

The authors have declared no competing interest.

