## supplemental for "CD96-mediated internalisation of ligand CD155 as a novel mechanism for immune regulation"

**Supplementary table S1: Antibodies**

| Target | Clone | Fluorophore | Company | Application |
|---|---|---|---|---|
| Anti-human Fc | M1310G905 | AlexaFluor647 | BioLegend | Flow cytometry |
| CD4 | SK3 | BUV563 | BD | Flow cytometry |
| CD155 | SKII.4 | BV421 | BD | Flow cytometry |
| CD96 | 6F9 | BV711 | BD | Flow cytometry |
| Anti-mCherry | 16D7 | AlexaFluor647 | ThermoFisher | Flow cytometry |
| TIGIT | 741182 | BUV395 | BD | Flow cytometry |
| CD226 | 11A8 | PE-Cy7 | BioLegend | Flow cytometry |
| CD96 extracellular domain (ECD) | BLR065G | Unconjugated | Bethyl Laboratories | WB |
| CD155 | D8A5G | Unconjugated | CST | WB |
| LAMP1 | D2D11 | Unconjugated | CST | WB, confocal, PLA |
| LAMP2 | H5B4 | Unconjugated | SantaCruz | WB |
| ATG5 | D5F5U | Unconjugated | CST | WB |
| ATG7 | F4V2N | Unconjugated | CST | WB |
| p62 | D5E2 | Unconjugated | CST | WB, confocal |
| LC3A/B | D3U4C | Unconjugated | CST | WB, confocal |
| RAB5 | C8B1 | Unconjugated | CST | Confocal |
| Anti-Human Fc | M1310G05 | Unconjugated | BioLegend | Confocal |
| Anti-rabbit | - | AF555 | ThermoFisher | Confocal |
| Anti-rat | - | AF647 | BioLegend | Confocal |
| Anti-GFP | 7.1, 13.1 | Unconjugated | Merck | Confocal, PLA |
| Beta-actin | 13E5 | Unconjugated | CST | WB |
| Alpha-tubulin | polyclonal | Unconjugated | CST | WB |
| Anti-rabbit | - | IRDye 800CW, IRDye 680RD | LI-COR | WB |
| Anti-mouse | - | RDye 800CW, IRDye 680RD | LI-COR | WB |

**Supplementary Table S2: Buffers**

| Buffer | Final concentration | Reagent | pH | Company |
|---|---|---|---|---|
| RIPA (lysis) | 50mM | Tris-HCl | 7.4 | ThermoFisher |
| | 5mM | EDTA | - | ThermoFisher |
| | 150mM | NaCl | - | Fisher |
| | 1% | Triton X-100 | - | Merck |
| | 10mM | Sodium orthovanadate ($Na_3VO_4$) | - | ThermoFisher |
| | 2mM | Sodium fluoride (NaF) | - | ThermoFisher |
| | 50mM | N-ethylmaleimide (NEM) | - | Fluorchem Ltd |
| | 1x | EDTA-free complete protease cocktail inhibitor | - | Merck |
| IP (wash) | 50mM | Tris-HCl | 7.4 | ThermoFisher |
| | 5mM | EDTA | - | ThermoFisher |
| | 150mM | NaCl | - | Fisher |
| | 1% | Triton X-100 | - | Merck |
| WB transfer | 1x | Tris-Glycine | - | Bio-Rad |
| | 5% | methanol | - | Fisher |
| WB staining | 1x | LI-COR Intercept Blocking Buffer | - | LI-COR |
| | 0.1% | Tween-20 | - | Merck |
| WB (wash), TBST | 1x | Tris-buffered saline | - | CST |
| | 0.1% | Tween-20 | - | Merck |

### Supp 1

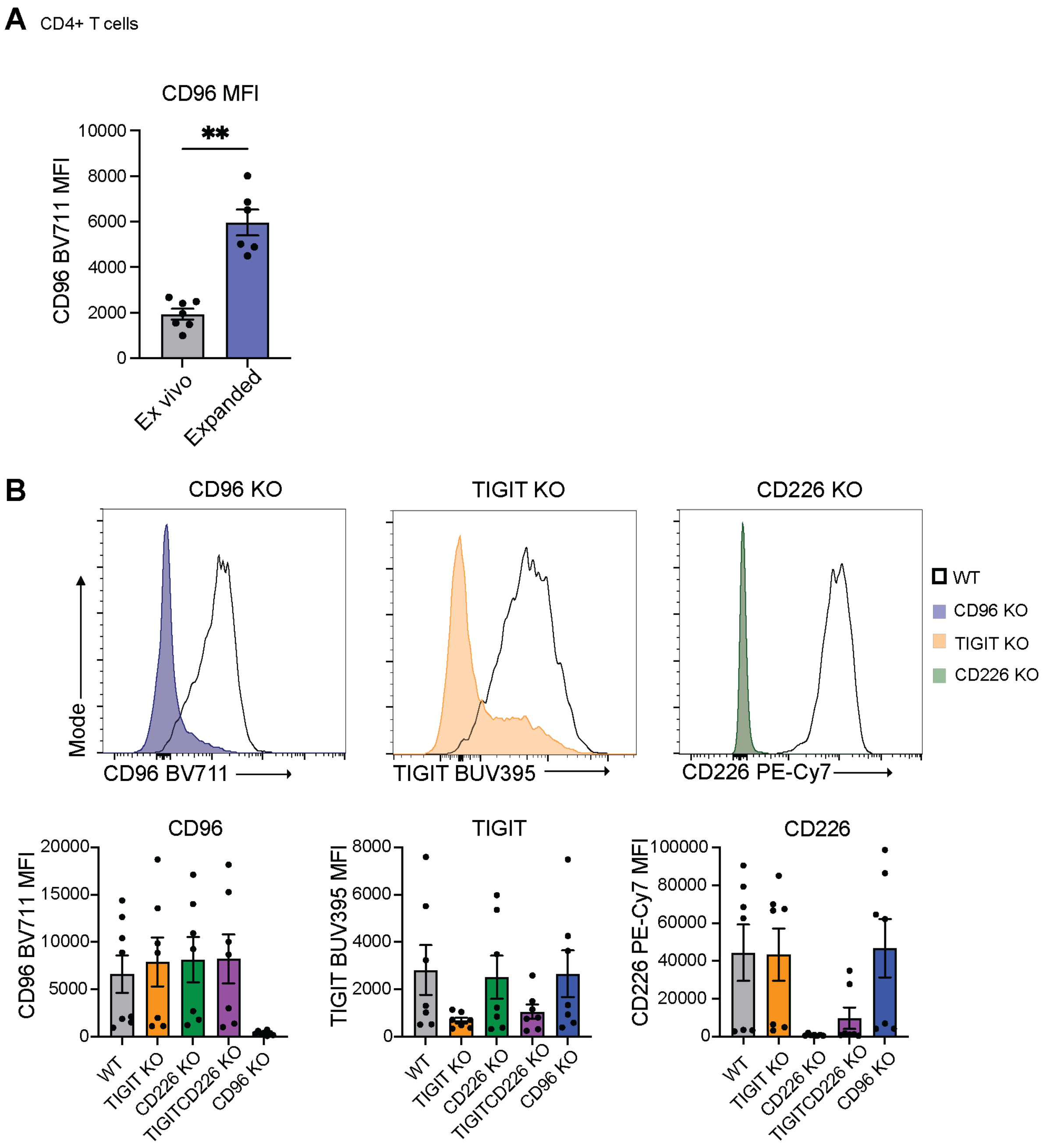


**Supplementary Figure S1**. **(A)** anti-CD96 BV711 MFI in ex vivo and expanded CD4+ T cells (n=7); Mann-Whitney test. **(B)** WT or CRISPR/cas9-mediated knockout (KO) of CD96, TIGIT, CD226, or TIGIT and CD226 double KO, expanded, primary human CD4+ T cells. Representative histograms and MFI of anti-CD96 BV711, anti-TIGIT BUV395, and anti-CD226 PeCy7 on expanded WT or CRISPRed CD4+ T cells, regulatory T cells (Tregs) and conventional T cells (Tconvs) (n=8). Data as mean±SEM; **p<0.01.

### Supp 2

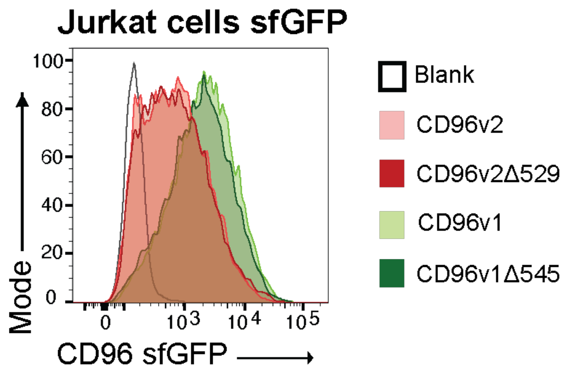


**Supplementary Figure S2**. Engineered Jurkat cells expressing WT CD96v1/v2sfGFP, or with cytoplasmic domain deletion (CD96v1Δ545/v2Δ529sfGFP); Representative histogram of (CD96)sfGFP expression for GFP-matched phenotype for WT vs cytoplasmic domain deletion within v1/v2 cell lines.

### Supp 3

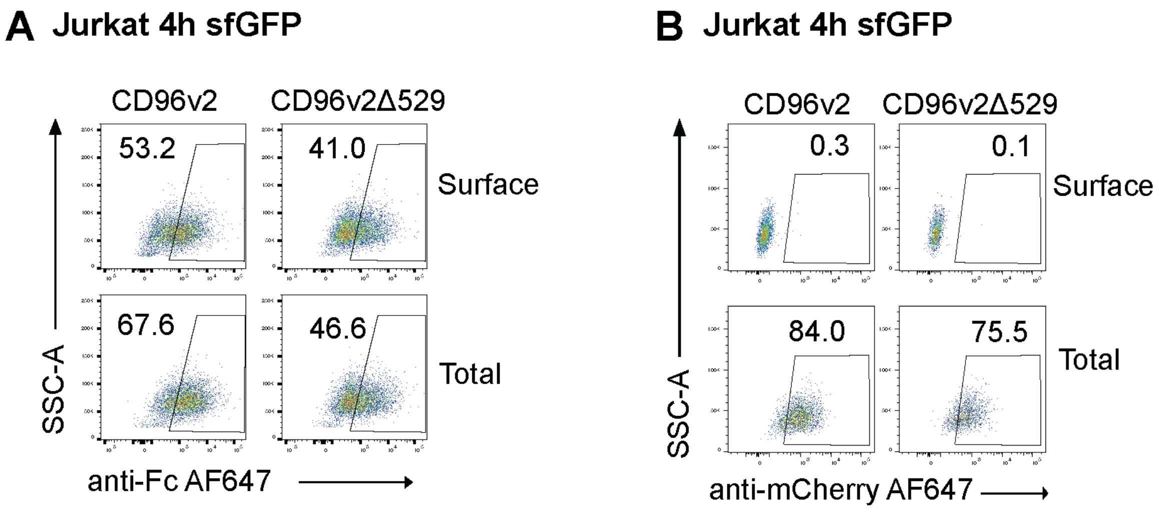


**Supplementary Figure S3**. **(A-B)** engineered Jurkat cells expressing WT CD96v2sfGFP or CD96v2Δ529sfGFP were incubated with soluble CD155Fc (5µg/mL, **A**) or 1:1 with DG75.CD155mCherry (**B**) over time at 37°C and 5% $CO_2$, and stained with anti-human Fc or anti-mCherry AF647 antibody on the surface or post-fixation/permeabilisation for the total stain; Representative plots for 4h surface and total anti-human Fc AF647 (**A**) or anti-mCherry AF647 (**B**) stain on sfGFP+ gated cells.
